# The Receptor Kinase BRI1 promotes cell proliferation in Arabidopsis by phosphorylation- mediated inhibition of the growth repressing peptidase DA1

**DOI:** 10.1101/2020.05.15.098178

**Authors:** Hui Dong, Caroline Smith, Rachel Prior, Ross Carter, Jack Dumenil, Gerhard Saalbach, Neil McKenzie, Michael Bevan

**Affiliations:** Dept of Cell and Developmental Biology, John Innes Centre, Norwich Research Park, Norwich NR4 UH, UK; Dept College of Agriculture, Nanjing Agricultural University, Nanjing, Peoples’ Republic of China 210095; Biological Chemistry Dept, John Innes Centre, Norwich Research Park, Norwich NR4 UH, UK; Sainsbury Laboratory Cambridge University, Bateman St, Cambridge CB2 1LR, UK

**Keywords:** Brassinosteroid, N-end rule protein stability, DA1 peptidase, Arabidopsis, cell proliferation, organ growth

## Abstract

Brassinosteroids (BR) have centrally important functions in plant growth by promoting cell proliferation and cell expansion through phosphorylation-mediated regulatory cascades that are initiated by perception of BR by receptor-like kinases of the BRI1 family. These BR-mediated growth responses have been explained by transcriptional controls mediated by phosphorylation of BES1/BZR1 transcription factors. Here we link BRI1-mediated phosphorylation to another growth regulatory network that directly mediates protein stability. BRI1 and its co-receptor BAK1 phosphorylate and inhibit the activities of the growth repressor DA1 by promoting the formation of high molecular weight complexes. Phospho-mimic forms of DA1 are less active while phospho-dead mutants have increased growth-repressive activity, while their regulatory monoubiquitylation is unaffected. BR inhibition of DA1 activity maintains higher levels of DA1 substrates such as UBP15 that sustain the potential for cell proliferation during leaf growth. Reduced BR levels lead to activation of DA1 peptidase activity and a transition from cell proliferation to cell growth and differentiation. This dual monoubiquitylation-phosphorylation regulation supports the key role of BR levels in maintaining the proliferative potential of cells during organ growth.

## Introduction

The final sizes and shapes of organs are determined by proliferation of small primordia of undifferentiated cells followed by cell growth and differentiation. Studies of organ growth in plants, in which the complexities of cell migration and death during organ growth are limited, have revealed spatial and temporal separation of primary cell divisions to the proximal region of young leaves that are progressively limited until cell growth drives primary leaf morphogenesis [1–3]. Combinations of modelling and imaging are developing a more integrated understanding of these spatio-temporal patterns of cell proliferation and growth in Arabidopsis leaves [4–6]. These studies show the final shapes of organs such as leaves are determined by anisotropic cell growth patterns and the final sizes of organs are determined by the duration of cell proliferation and the extents of cell growth. Proposed regulators, oriented by polarity fields, influence the rate and directions of cell growth, the competence of cells to divide, and size thresholds of division [5].

The transition from cell proliferation to cell growth and differentiation is a critical determinant of final organ size [7]. This involves a progressive reduction in the capacity of proximal zones of cells to proliferate [8] and involves large-scale transcriptional changes [2,7]. Apical-basal gradients of miRNAs396 and 319 provide one mechanism to limit the activities of transcription factors that promote cell proliferation to proximal regions of developing leaves. These include members of the Growth Regulatory Factor (GRF) family [9,10], which are recognised by miRNA396, and members of the Teosinte-Branched1-Cycloidea-PCF (TCP) family of transcription factors, which are recognised by miRNA319a(JAW) [11]. These transcription factors and others (reviewed in [12]) contribute to the maintenance of mitotic cycles and the inhibition of endoreduplication, cell growth and differentiation. The proteome is also regulated as part of the transition from cell proliferation to endoreduplication and cell growth during organ formation. Members of the DA1 peptidase family proteolytically cleave and inactivate, *via* N-end rule mediated proteolysis, several different proteins involved in the transition from cell proliferation to endoreduplication [13,14]. DA1 substrates include TCP15 and TCP22, which promote cell proliferation and repress endoreduplication, and UBP15 [15] that promotes cell proliferation. DA1-mediated cleavage of these proteins is thought to contribute to the cessation of cell proliferation and the initiation of endoreduplication and cell growth, thus limiting organ sizes. DA1 family peptidase activity is activated by multiple monoubiquitylation mediated by the E3 ligases Big Brother (BB) and DA2 [14], and this is reversed by de-ubiquitylation mediated by UBP12 and UBP13 [16]. Reduced UBP12 and UBP13 activities lead to smaller leaves with fewer smaller cells, consistent with their role in limiting the activities of DA1 family growth repression. The timing and spatial location of this regulatory ubiquitylation and de-ubiquitylation are currently not clear, but it seems likely that the activities of DA1 and other repressors of cell proliferation need to be limited in order to establish and maintain patterns of cell proliferation.

Plant growth regulators have centrally important roles in organ growth. Mutants deficient in brassinosteroid (BR) perception or biosynthesis have small organs due to reduced cell numbers and cell sizes. Complementation of BR signalling defects in the epidermal cell layer rescued normal leaf growth [17], while depletion of BRs in the epidermis inhibited leaf growth, suggesting leaf growth depends on local BR synthesis and perception in the epidermis. Detailed analyses of leaf growth in a BR-deficient mutant demonstrated that low levels of BR lead to reduced cell size and cell numbers that accounted for reduced leaf areas [18]. This provided further support for key roles of BR biosynthesis and signal perception in maintaining a balance between cell proliferation and cell growth during leaf development. A BR-dependent mechanism also balances patterns of cell proliferation and quiescence in the root tip ([19,20] and BR perception in protophloem tissues promotes cell proliferation in the root meristem [21], showing non-cell autonomous effects of BR perception on organ growth. BR levels are reduced at leaf margins by LOB transcription factors that activate *BAS1* encoding a BR-inactivating enzyme [22]. Low levels of marginal BR permits expression of boundary-specifying CUC genes, while CUC genes are repressed in meristematic cells by higher levels of BR that promote BZR1-mediated transcriptional repression [22,23]. Together these observations indicate an important role for BR levels and signalling in promoting and maintaining zones of cell proliferation in developing leaves.

Here we show that DA1 is a substrate of the kinase activities of the brassinosteroid receptor BRI1 and its co-receptor BAK1. Phosphorylation of DA1 by BRI1 and BAK1 leads to reduction of its peptidase activity, while not affecting regulatory monoubiquitylation. Non-phosphorylated forms of DA1 retain peptidase activity and are more active growth repressors while phospho-mimic forms of DA1 are less active growth repressors. Consistent with this, DA1 peptidase activity is reduced by BR treatment and levels of DA1 peptidase substrates increase in response to BR. These observations are consistent with the roles of BR-mediated signalling in promoting organ growth and establish a link between BR-mediated signalling and the regulation of protein stability during organ growth.

## Results

### DA1 influences the extent of proliferative zones during early stages of leaf growth

Segmentation and computational analyses were used to quantify the influence of DA1 on the numbers, sizes and distributions of cell sizes across whole leaves and their contributions to growth. An epidermal cell membrane localised marker generated by a *pAtML1::mCitrine-RC12A* transgene (pAR169) in Col-0 and *da1-1* genotypes were imaged and segmented in the first leaf between 7-11 das (days after stratification) in the abaxial epidermal layer (Figure S1). Data on the location and area of each segmented epidermal cell was generated and relative cell densities calculated. At the earliest observed time of 7 das, *da1-1* lines had significantly increased cell numbers compared to Col-0, and these steadily increased until the *da1-1* had over twice the number of cells than Col-0 at 11 das (Figure 1A). Figure 1B shows that segmented cell areas varied widely at all time points, with a tendency for *da1-1* cells to be smaller. Final epidermal cell sizes were smaller in *da1-1* compared to Col-0 (Figure 1B), showing that increased leaf growth in *da1-1* can be accounted for by increased cell numbers. We used cell density as a proxy for defining zones of cells with the potential to proliferate, as cells can only increase in size once they exit mitosis, endoreduplicate and differentiate [12]. This increase in cell area reduces the density of cells. We used a conservative value of 80% of the maximum cellular density to delineate regions of higher potential to proliferate. Cell densities were calculated and are shown in heatmaps in Figure 1C and graphically in Figure 1D. At the first two time-points, zones of high cell density in *da1-1* accounted for a larger proportion of leaf area than in Col-0. At each time-point the proportion of this area progressively reduced in both *da1-1* and Col-0 due to cell growth. This demonstrated that reduced DA1 growth repression in the *da1-1* mutant increased the initial numbers of cells with the potential to proliferate and maintained this larger number during early stages of leaf growth. This larger number of cells accounted for increased organ size.

**Figure 1.**
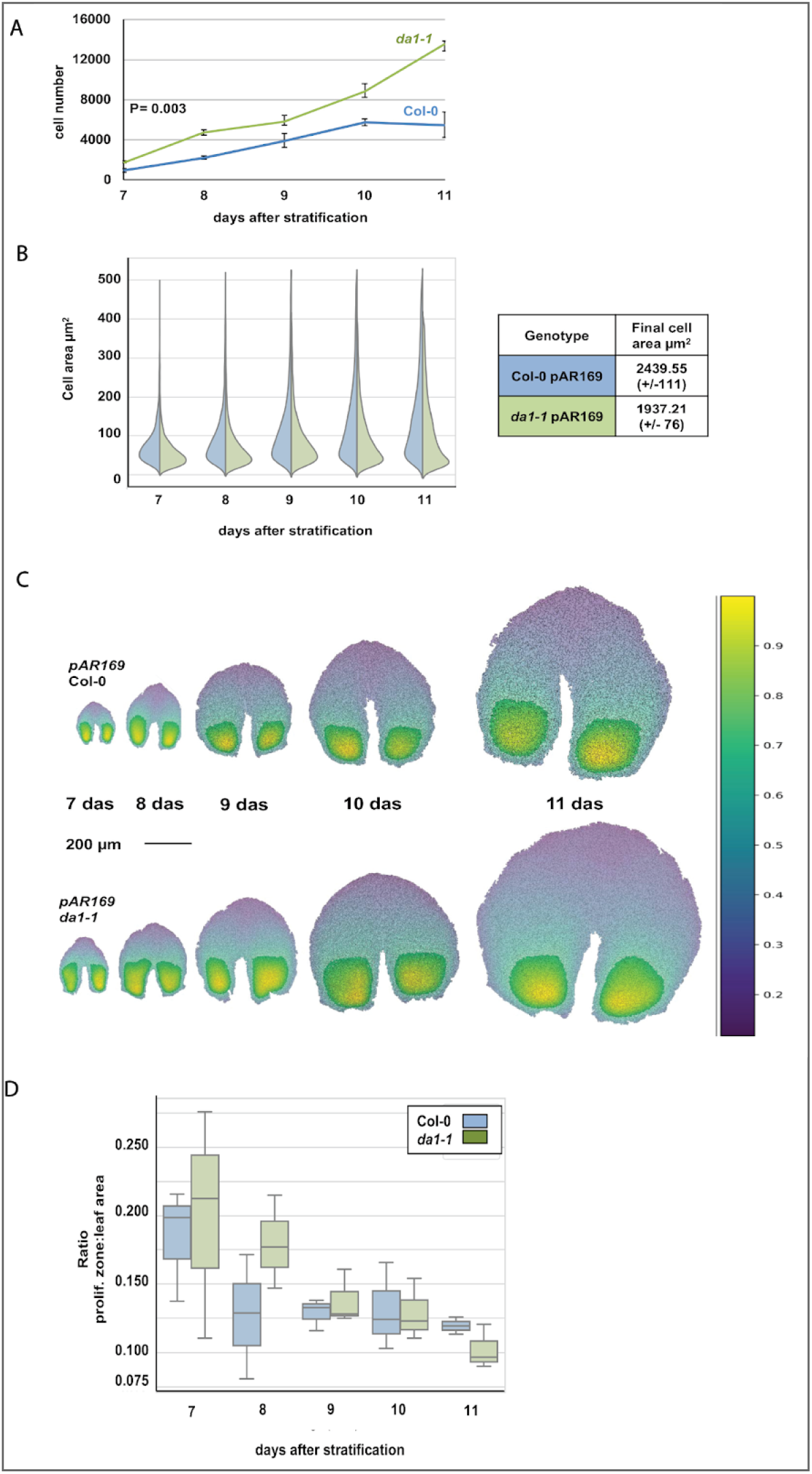
Cellular analysis of leaf growth in *da1-1* mutants. (A) Graph of mean abaxial segmented cell numbers between 7-11 days after stratification (das) in Col-0 and da1-1 pAR169 seedlings. At the earliest imaged timepoint the numbers of cells were significantly increased in *da1-1* pAR169. Data are given as mean ± SEM. n= first leaves of 3 independent seedlings. P values were determined by Student’s t-test. (B) Graph of mean abaxial segmented cell areas during leaf growth in Col 0 and *da1-1* pAR169 seedlings. The top and bottom 5% values were trimmed. Cell areas (between 1200-3000 from each of 3 independent seedlings) were measured. The inset Table shows final cell sizes in the abaxial epidermis after 35 days growth determined by SEM. n= 20 cells per genotype. (C) The heat maps show the distribution of zones of relatively high cell density that reflect the potential to proliferate between 7-11 das in Col 0 and *da1-1* pAR169 leaves. (D) Graph showing the ratio of areas of cells with density >0.8 divided by the total area of all cells between 7-11 das.

### Genetic and physical interactions of BRI1 and DA1/DAR1

To identify mechanisms that may modulate the growth repressive activity of DA1 peptidase a yeast-two-hybrid (Y-2-H) screen for potential interacting proteins was performed. We consistently identified the cytoplasmic domain of the Receptor-Like Kinase (RLK) TMK4 [24–26] among the interactors. (Figure S2). We therefore surveyed genetic and physical interactions of DA1 with a range of RLKs with roles in cell proliferation and growth. Among several genetic and physical interactions of DA1 and DAR1 with Receptor-Like Kinases (RLKs), we describe those with the RLK BRI1, a receptor for brassinosteroid growth regulators [27].

The large cotyledon areas of *da1dar1* double loss-of-function mutants [14] (which were used as loss-of-function alleles compared to the interfering allele *da1-1* in order to facilitate interpretation of genetic interactions) were decreased to near those of wild-type by the temperature-sensitive *bri1-301* mutant in *da1dar1bri1-301* lines at the non-permissive temperature 29°C (Figure 2A and B). This allele of *BRI1* has reduced kinase activity and protein levels at non-permissive temperatures [28] and was used for ease of creating and maintaining mutant combinations. Light-grown hypocotyl lengths were also increased in the triple mutant compared to *bri1-301* (Figure 2A). Cell areas in *da1dar1bri-301* were the same as in *bri1-301*, consistent with increased cell numbers leading to increased cotyledon areas (Figure 2C). Hypocotyl elongation in the dark (commonly used to assess BR action) showed the same patterns of relief of the short hypocotyl phenotype of *bri1-301* by the *da1dar1* double mutant (Figure 2D and 2F), with cell lengths being quite similar in both (Figure 2G). Brassinazole (BRZ) treatment to reduce endogenous BR levels further reduced hypocotyl lengths in *bri1-301*, showing its partial suppression of BR-mediated growth responses at 29°C (Figure 2E and 2F). The *da1dar1bri1-301* mutant had significantly longer hypocotyls than *bri1-301* in these conditions. This was consistent with analyses of light-grown plants (Figure 2A), showing that BR-mediated responses were elevated by loss of DA1/DAR1 function. The canonical BR signalling pathway regulating BZR1-mediated transcription is intact in *da1dar1* lines, shown by similar levels of induction of the BZR1 target genes *EXP8* and *SAUR AC1* [29] in response to epi-Brassinolide (BL) in both Col-0 and *da1dar1*, and the absence of BL-mediated induction in *bri1-301* in these conditions (Figure 2H and 2I). Taken together, these observations show that suppression of growth by DA1 and DAR1 is reduced by BR-mediated signalling through BRI1 in a pathway that appears to be independent from the canonical BR signalling pathway that regulates gene expression.

**Figure 2.**
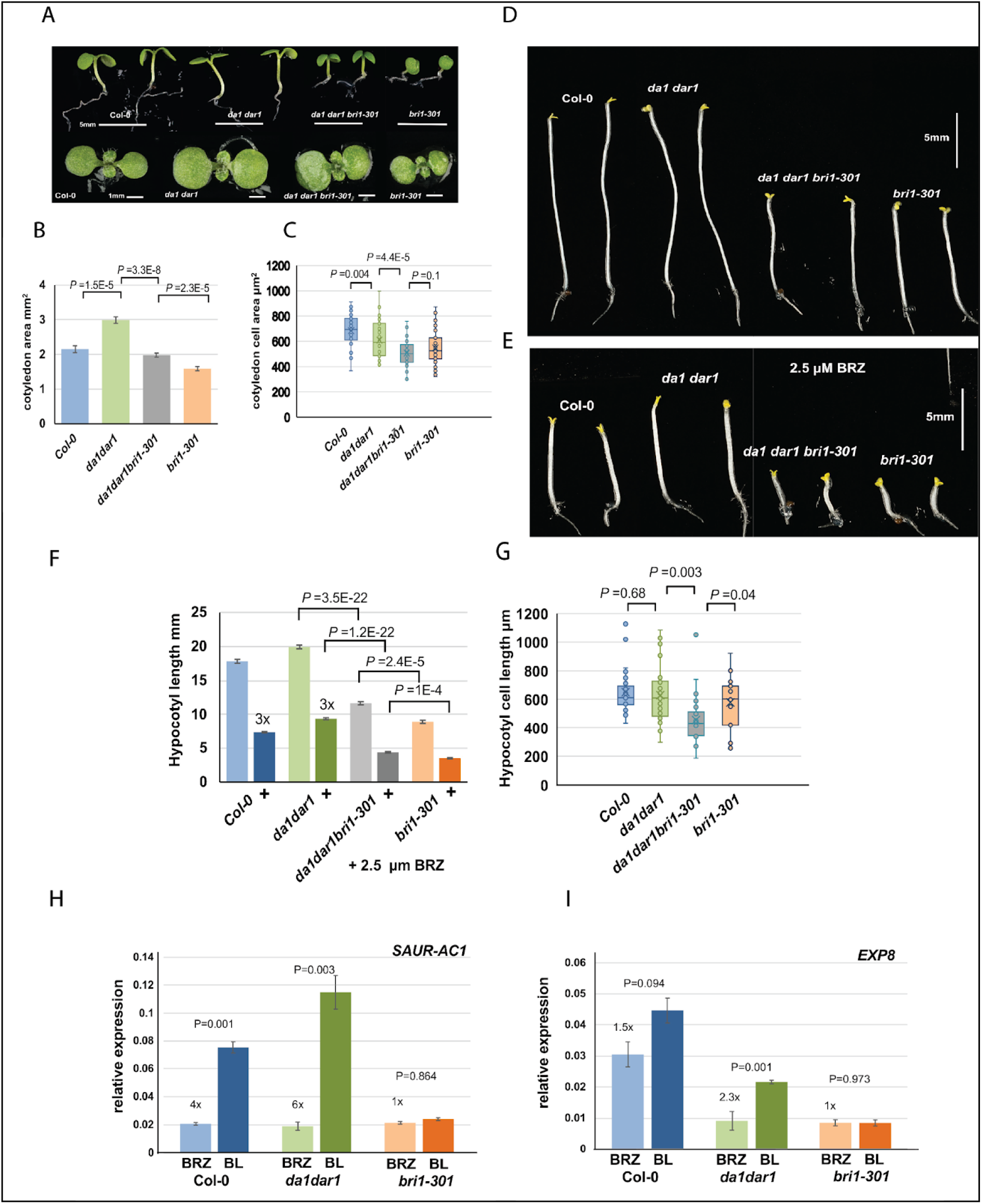
Genetic interactions of DA1 with BRI1. (A) Seedling (upper panel) and cotyledons (lower panel) of 7 day old Col-O, *da1dar1*, *da1dar1bri1-301* and *bri1-301* seedlings grown at 29°C. (B) Cotyledon areas (n=25-30) and cell areas (C), n = 10-15) of 7 day old Col-O, *da1dar1*, *da1dar1bri1-301* and *bri1-301* seedlings. Data are given as mean ± SEM. P values were determined by Student’s *t*-test. (D) Hypocotyl elongation in response to DMSO in Col-0, *da1dar1, da1dar1bri1-301, bri1-301* 5 day old dark grown seedlings. (E) Hypocotyl elongation in response to 2.5 μM BRZ in DMSO in Col-0, *da1dar1, da1dar1bri1-301, bri1-301* 5 day old seedlings (n=20-25) grown on vertical plates in the dark. (F) Hypocotyl (n=20-25) and hypocotyl cell lengths (G) (n=10-15) of Col-O, *da1dar1*, *da1dar1bri1-301* and *bri1-301* 5 day old dark-grown seedlings. Data are given as mean ± SEM. P values were determined by Student’s *t*-test. (H) *EXP8* and (I) *SAUR AC1* expression in seedlings in response to 2.5 μM BRZ or 1.0 μM BL in Col-0, *da1dar1* and *bri1-301* seedlings. All seedlings were grown at 29°C for 7 days after stratification. Expression levels are relative to *EF1 ALPHA* expression. The fold-difference in expression between BRZ and BL treatment is shown. Data are given as mean ± SEM. n = 3 biological replicates. P values were determined by Student’s *t*-test.

These genetic interactions between loss-of-function DA1 and DAR1 mutations and a reduced BRI1 function mutant suggested they may function together. Figure 3A shows that BRI1-GFP and DA1-mCherry co-localised at the plasma membrane of transfected *da1dar1* mesophyll protoplasts. DA1-mCherry was also located in the cytoplasm. Fractionated plasma membrane samples from transgenic lines expressing DA1-GFP that complemented *da1dar1* (see Figure 5C for complementation data) were enriched for DA1-GFP, consistent with colocalization data (Figure 3B). These experiments reveal a predominant location of DA1 at the plasma membrane, consistent with a potential role in BRI1-mediated signalling mechanisms. DA1-GFP interacted directly with BRI1-3FLAG and more weakly with BAK1-3FLAG fusion proteins when co-expressed in *N. benthamiana* leaves (Figure 3C). In transgenic *da1dar1* Arabidopsis plants expressing *35S::HA-DA1* and *BRI1::BRI1-GFP* fusion proteins (Figure 3D) this interaction was also observed. These data indicate that DA1 fusion proteins can co-locate and physically interact with BRI1 fusion proteins, suggesting a potential direct role in BRI1-mediated signalling processes.

**Figure 3.**
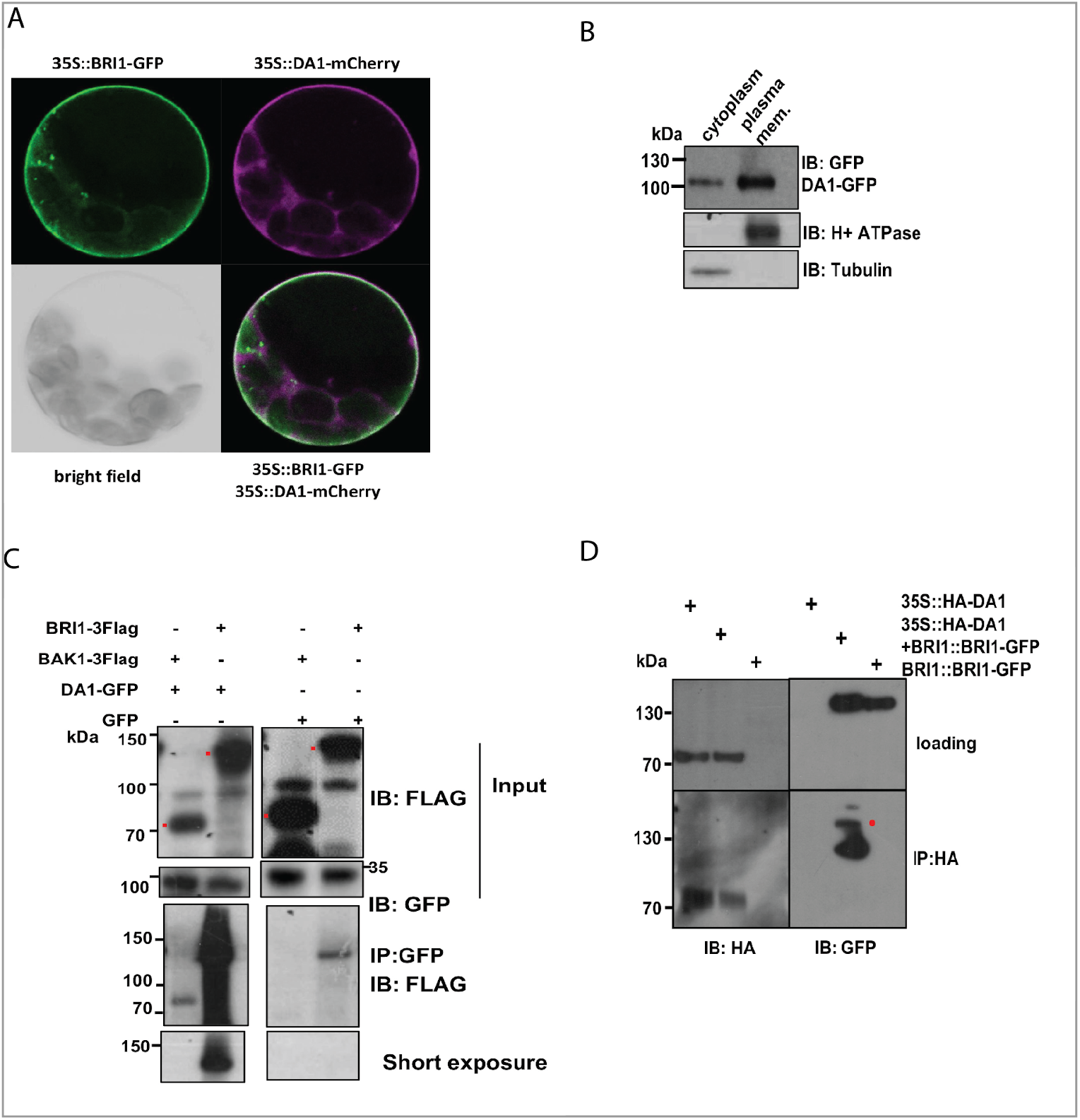
Co-location and physical interaction of DA1 and BRI1. (A) Co-location of DA1-mCherry and BRI1-eGFP on the plasma membrane of transfected mesophyll protoplasts. (B) Immunoblots of plasma membrane and cytoplasmic fractions from transgenic DA1::DA1-GFP seedlings. (C) Immunoprecipitation of BRI1-3FLAG and BAK1-3FLAG by DA1-GFP in transfected *Nicotiana benthamiana* leaves. The red dots indicate BRI1-3FLAG and BAK1-3FLAG protein. (D) Immunoprecipitation of BRI1-GFP by HA-DA1 in *35S::HA-DA1 BRI1::BRI1-GFP* transgenic *da1dar1* seedlings. The red dot indicates full-length BRI1-GFP protein.

### BRI1 and BAK1 phosphorylate DA1

DA1 is a peptidase that cleaves a variety of growth regulatory proteins, promoting their degradation and limiting cell proliferation [14], therefore we tested whether BRI1 and BAK1 were substrates. Neither of their cytoplasmic C-terminal domains tested were cleaved by DA1 in Arabidopsis protoplasts (Figure S3), suggesting that BRI1 and BAK1 might instead influence DA1 activity. These proteins directly phosphorylate each other and BRI1 phosphorylates the BR-signalling kinases BSK2 and BSK3 [14,30] and the BRI1 inactivating protein BKI1 [31]. To test if BRI1 and BAK1 directly phosphorylate DA1, cytoplasmic domains (cd) of BRI1 or BAK1 (and their kinase dead mutants) were fused to -FLAG were co-expressed with GST-DA1 in *E.coli*. Mass spectrometric analysis of affinity purified GST-DA1 identified multiple BRI1- and BAK1-dependent phosphoserine and phosphothreonine residues on DA1 (and Figure 4A and Figure S4). We focussed on two clusters of phospho-sites in functionally characterised regions of DA1 that were consistently identified in independent mass spectrometry runs: T90, S91 and S97 were close to the ubiquitin interaction motifs, and S363, T367, S369 and T370 were in a conserved region of DA1 (see inset to Figure 4A) that harbours monoubiquitylation sites, the *da1-1* interfering allele (R358K) and its suppressor *sod1-2* L362F allele [13]. Mutation of DA1 T90, S91 and S97 to either phospho-mimic D or phospho-dead A amino acids had no effect on cleavage of DA1 substrates BB-FLAG or FLAG-UBP15 (Figure S5). We therefore focussed on characterising the potential roles of phosphorylation of T367, S369 and T370 as the most consistently abundant phospho-forms determined by mass spectrometry. First, BRI1-cd and BAK1-cd phosphorylated DA1 *in vitro*, while kinase-dead versions of BRI1-cd and BAK1-cd were not active on DA1 (Figure 4B). DAR1 was also phosphorylated by BRI1-cd and BAK1-cd at lower levels than those observed for DA1 (Figure 4B lower panels). Mutation of T367, S369 and T370 to non-phosphorylatable alanine residues (T367A, S369A, T370A; termed DA1-3A and T385A, T387A, T388A, termed DAR1-3A) abolished BRI1-cd and BAK1-cd mediated phosphorylation, demonstrating that this trio of adjacent T and S amino acids on DA1 and DAR1 were major sites of phosphorylation by BAK1 and BRI1 *in vitro*. A transgenic line expressing HA-DA1 from the 35S promoter that complemented the *da1dar1* mutation (Figure S6A) was used to analyse *in vivo* phosphorylation using 2D gel electrophoresis. BL treatment led to a higher proportion of acidic forms of HA-DA1 compared treatment with BRZ alone (Figure 4C), consistent with increased phosphorylation of HA-DA1 in response to BL-mediated stimulation of DA1 phosphorylation. Transgenic lines expressing either HA-DA-3A phospho-dead or phospho-mimic forms (T367D, S369D, T370D, termed HA-DA1-3D) (Figure S6B) did not show these changes to lower pI forms after similar treatments with BL and BRZ. These experiments showed that HA-DA1 is a direct substrate of BRI1 and BAK1 and that BL treatment leads to reduction of HA-DA1 pI that is consistent with *in vivo* phosphorylation of HA-DA1 by BRI1 in response to BL treatment.

**Figure 4.**
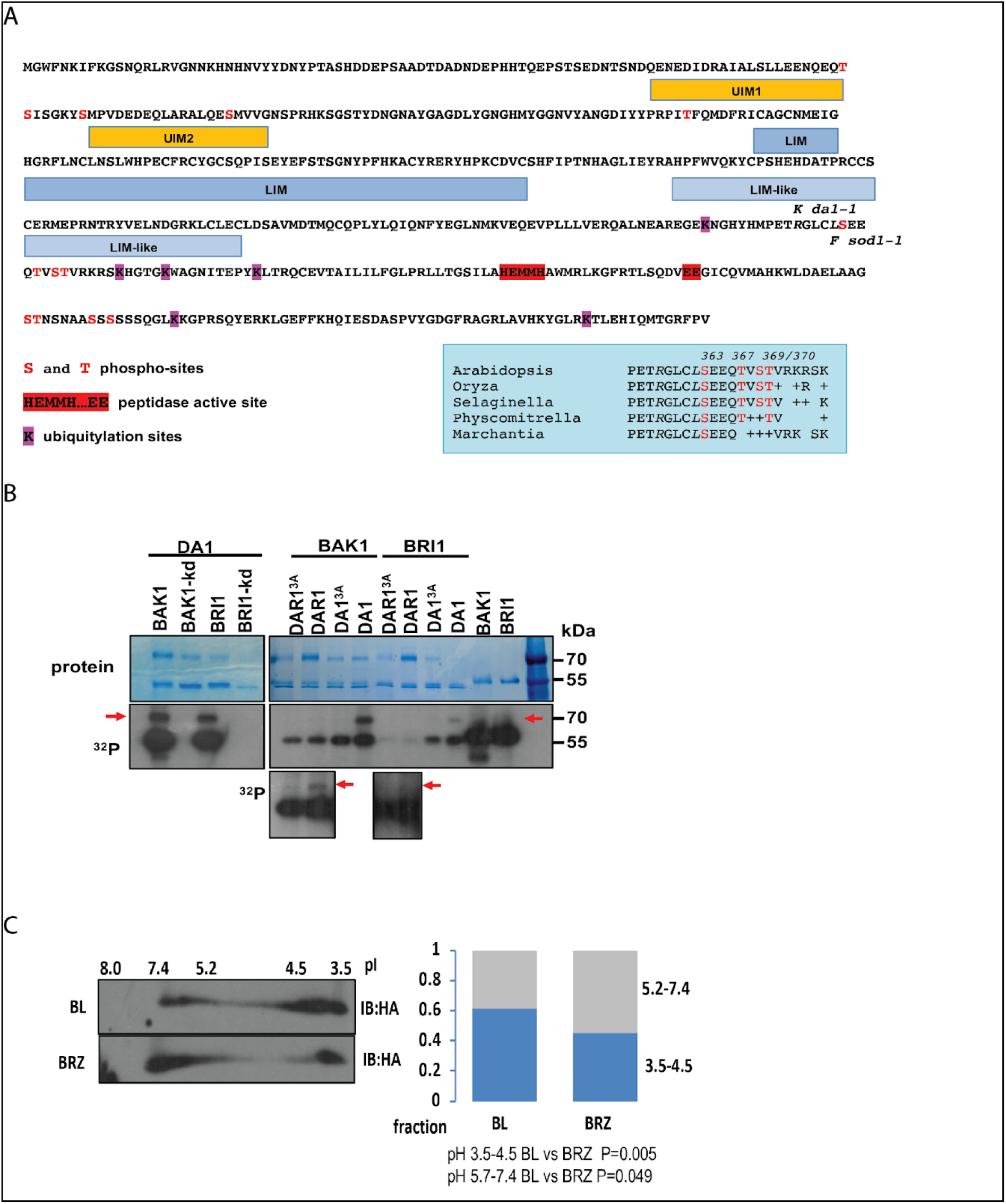
BRI1 and BAK1 phosphorylate DA1. (A) Location and conservation of phospho-sites on the DA1 protein sequence determined by co-expression of BRI1 and DA1-GFP in *E.coli* and mass spectrometry. The locations of known functional and structural motifs are shown for reference. The inset shows the phospho-sites and their conservation. UIM Ubiquitin Interaction Motif; LIM - Lin11-Isl-1-Mec-3 domain. (B) *In vitro* phosphorylation of DA1 and DAR1 by BRI1 and BAK1. The upper gel shows protein levels and the lower panel shows autoradiographic detection of ^32^P-labelled proteins. A longer exposure in the lower panel shows DAR1 phosphorylation. The red arrows indicate phosphorylated forms of DA1 and DAR1. The mobilities of BAK1 and BRI1 were slightly increased due to the additional loading of DA1 and DAR1. (C) Immunoblot of two-dimensional gels of HA-DA1 from transgenic seedlings treated with epi-brassinolide (BL) or brassinozole (BRZ) for 2 hrs. The right panel shows the proportion of HA-DA1 in acidic and neutral regions of the gels in response to BL and BRZ (n=5 independent seedling microsome extractions). P values were determined by Student’s *t*-test.

### Phosphorylation influences the peptidase activity and homo-dimerization of DA1

In order to understand how phosphorylation might influence DA1 activity we examined the peptidase activities of phospho-dead HA-DA1-3A and phospho-mimic HA-DA1-3D on its substrate BB [32]. Figure 5A shows that HA-DA1-3D had undetectable peptidase activity on BB-FLAG after co-expression in *da1dar1* protoplasts, whereas the HA-DA1-3A phospho-dead form exhibited the same levels of peptidase activity as HA-DA1. As the peptidase activity of DA1 is activated by multiple monoubiquitylation [14], we tested whether FLAG-DA1-3D was ubiquitylated by the E3 ligase DA2. Figure 5B shows that FLAG-DA1-3D was ubiquitylated to the same levels as FLAG-DA1, suggesting that reduced peptidase activity of phospho-mimic DA1-3D cannot be explained by reduced mono-ubiquitylation. Therefore BRI1- and BAK1-mediated phosphorylation of DA1 reduces its peptidase activity independently of ubiquitin-mediated activation of DA1 activity, consistent with BL promoting growth by inhibiting the activity of a growth repressor (Figure 2). The influence of DA1-3A and DA1-3D mutations on plant growth were assessed by the extent of complementation of the large organ phenotype of the *da1dar1* double mutant. Figure 5C shows that transgenic lines expressing genomic constructs of DA1-GFP (including splice sites and 5’ and 3’ flanking regions) complemented DA1 function by restoring normal petal sizes. In contrast, DA1-3A-GFP genomic constructs had significantly smaller petals than Col-0 lines expressing wt DA1-GFP (Figure 5C), showing higher DA1 growth-repressive activity of non-phosphorylatable DA1. Transgenic lines expressing DA1-3D-GFP had a relatively weak complementing activity compared to DA1-GFP, indicating reduced DA1 growth-repressive activity. These observations were consistent with reduced peptidase activity of DA1-3D (Figure 5A), while the increased activity of DA1-3A indicated that BR-mediated phosphorylation may modulate ubiquitin-activated DA1 *in vivo*. This was tested using DA1 activity assays in transfected protoplasts treated with BL or BRZ. Figure 5D shows that HA-DA1 peptidase activity was modulated by BL levels, as treatment with BL reduced peptidase activities on both BB-FLAG and FLAG-UBP15 [15], while BRZ treatment increased DA1 peptidase activities. This modulation was not seen in HA-DA1-3D, which had no detectable peptidase activity towards either substrate in either BL- or BRZ-treated protoplasts as expected, while HA-DA1-3A had a high level of activity in protoplasts treated with both BL and BRZ. This supported our interpretation of the complementation assays by showing that BL promotes organ growth by phosphorylation of DA1 T367, S369 and T370 and reduction of its peptidase activities.

**Figure 5.**
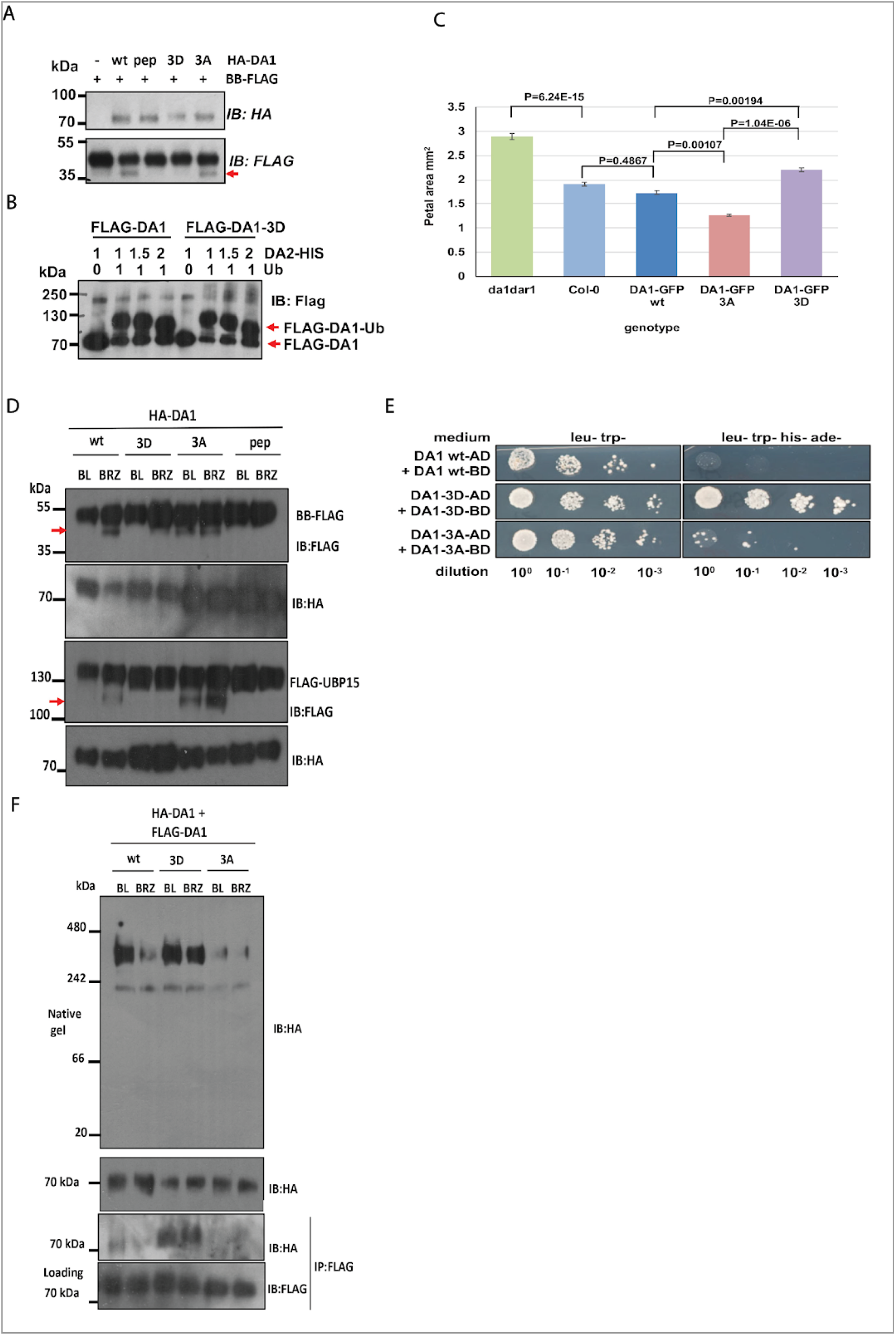
Functional analysis of DA1 phosphorylation. (A) Immunoblot showing peptidase activity of HA-DA1 wt, peptidase mutant, phospho-mimic 3D and phospho-dead 3A mutant proteins on the substrate BB-FLAG. The red arrow indicates the BB-FLAG cleavage product. HA-DA1 wt has peptidase activity, while HA-DA1 peptidase mutant has a mutation of the active site that abolishes activities on BB and other substrates. (B) Immunoblot showing ubiquitylation of HA-DA1 and HA-DA1 phospho-mimic mutant 3D by the E3 ligase DA2. The red arrow indicates ubiquitylated forms of HA-DA1 and HA-DA1-3D. Ub is Ubiquitin. (C) Complementation of *da1dar1* seedlings by *DA1::DA1 wt-GFP, −3D-GFP* and *-3A-GFP* constructs measured by petal areas (n=5 petals from each of 10 independent transgenic lines for each construct). Data are shown as means of means (n = 10) ±SEM. P values were determined by Student’s *t*-test. (D) Cleavage of BB-FLAG and FLAG-UBP15 by HA-DA1 wt, HA-DA1-3D phospho-mimic, HA-DA1-3A phospho-dead and peptidase mutant proteins in transfected *da1dar1* mesophyll protoplasts treated with BRZ or BL. The red arrows indicate BB-FLAG and FLAG-UBP15 cleavage products. (E) Yeast-2-hybrid analysis of interactions between DA1 wt, DA1-3D phospho-mimic and DA1-3A phospho-dead mutants. (F) Interactions of HA-DA1 wt, HA-DA1-3D phospho-mimic and HA-DA1-3A phospho-dead mutant proteins with DA1-FLAG wt, −3D and −3A proteins respectively in the same samples of proteins expressed in transfected *da1dar1* mesophyll protoplasts treated with BRZ or BL. The upper panel is an immunoblot of native protein samples on a 4-12% native polyacrylamide (PA) gel, the lower panels are immunoblots of denatured protein samples on 4-16% SDS-PA gels. These show, from the top, equal loading levels of HA-DA1 proteins on the native gel, HA-DA1 proteins immuno-purified through interaction with DA1-FLAG proteins, and immuno-purified DA1-FLAG proteins as the pull-down control.

We explored possible mechanisms that may account for this reduction in DA1 peptidase activity. In other studies we identified homo- and hetero-meric interactions between DA1 and other family members. Part of this yeast-2-hybrid (Y-2-H) dataset revealed DA1 homodimerization (Figure 5E). DA1:DA1 and DA1-3A:DA1-3A interactions were relatively weak, while DA1-3D:DA1-3D interactions were stronger. These patterns of interactions between DA1 proteins were also observed in Arabidopsis. Figure 5F shows co-expression of HA-DA1 and DA1-FLAG wt, −3D and −3A proteins in *da1dar1* mesophyll protoplasts. An immunoblot of a non-denaturing native polyacrylamide gel (upper panel) revealed forms of HA-DA1 migrating at approximately 300 kDa that were more abundant in protoplasts treated with BL compared to BRZ. HA-DA1-3D also migrated as similar high MW forms that were equally abundant in BL- and BRZ-treated protoplasts. Despite equal expression levels with HA-DA1-3D, high MW forms of HA-DA1-3A were much less abundant than those observed for wt HA-DA1 in BL-treated protoplasts and for HA-DA1-3D in both BL- and BRZ-treated protoplasts. In the same transfected protoplasts used for the native gel, pull-down experiments of HA-DA1, HA-DA1-3D and HA-DA1-3A by DA1-FLAG, DA1-3D-FLAG and DA1-3A-FLAG respectively showed increased interactions between HA-DA1-3D and DA1-3D-FLAG that were similar in both BLand BRZ-treated protoplasts. In contrast, interactions between wt DA1 proteins were increased by BL treatment, while no detectable interactions were observed between DA1-3A proteins. These data indicated that BL treatment increased interactions between DA1 molecules that lead to the formation of a high MW complex. BRZ treatment leads to lower levels of this complex, similar to the reduced levels of high MW complex and interactions observed between non-phosphorylatable and more active DA1-3A proteins. The stronger interactions of DA1-3D phospho-mimic proteins in both BL- and BRZ-treated protoplasts, and the increased levels of high MW complexes, is consistent with BL-mediated phosphorylation of T367, S369 and T370 leading to increased interactions between DA1 molecules. Taken together with the reduced peptidase activities of DA1-3D shown in Figure 5D, this suggested that BL-mediated phosphorylation of DA1 leads to the formation of HMW forms that are less enzymatically active.

### BL treatment stabilises DA1 substrates in transgenic plants

The modulation of DA1 activity by BL- and BRZ-treatment, reduced enzymatic activities of HA-DA1 phospho-mimic protein, and reduced growth restrictive activity of DA1-3D-GFP phospho-mimic protein, suggested that BL promotes growth in part by limiting the cleavage of DA1 peptidase substrates. DA1 peptidase cleavage destabilises its substrates BB and UBP15 [14,15] through the N-end rule pathway. Transgenic seedlings expressing complementing forms of UBP15-GFP and BB-Venus were treated with BRZ and the relative abundance of proteins determined after either further treatment with BRZ or with BL. Figure 6A shows levels of UBP15-GFP rapidly increased within 1 hr of BL treatment, contrasting to no detectable change in BRZ-treated seedlings over this time. Similar increases in BB-Venus levels were seen soon after BL treatment, while a non-cleaved mutant of BB (BB-AYGG) maintained similar levels between BRZ and BL treated seedlings (Figure 6B). These data showed that BL levels rapidly increase the abundance of DA1 substrates in transgenic plants.

**Figure 6.**
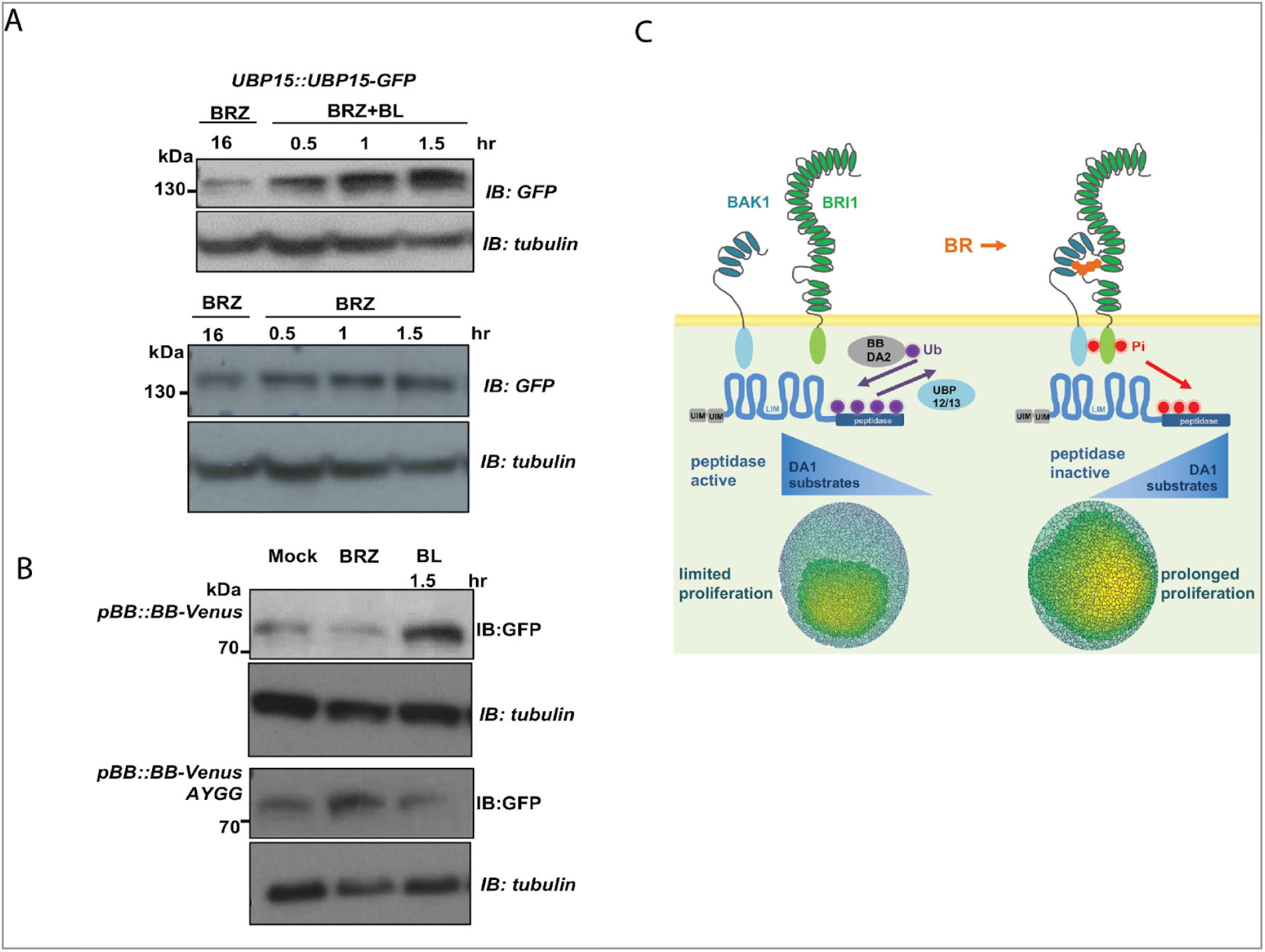
BL stabilises UBP15-GFP and BB-Venus in transgenic plants. (A) Immunoblot of UBP15-GFP protein levels in transgenic *UBP15::UBP15-GFP* 10-12 day old seedlings in response to 2.5 μM BRZ or 2.5 μM BRZ plus 1 μM BL treatments. Protein loading is shown by tubulin levels. (B) Immunoblot of BB-Venus protein levels in transgenic *pBB::BB-Venus* and the *pBB::BB-AYGG-Venus* mutant in response to either BRZ or BRZ plus BL treatment. The AYGG mutant of BB-Venus is not cleaved by DA1 and serves as a negative control. Protein loading is shown by tubulin levels. (C) The diagram summarises how BR regulates DA1 peptidase activity by BRI1- and BAK1-mediated phosphorylation. This reduces DA1 peptidase activity and increases levels of DA1 substrates [14] that promote cell proliferation. Low BR levels reduce DA1 phosphorylation and increase the activities of DA1 peptidase activation by monoubiquitylation. This reduces DA1 substrate levels and limits cell proliferation.

## Discussion

Brassinosteroids have well established roles in signalling from the cell surface to the nucleus, where they regulate the expression of multiple genes through BZR1 and BES1 transcription factors. Outputs of this signal transduction pathway include non-cell autonomous promotion of cell growth and division [33,34], and notably the promotion of cell proliferation during leaf growth [18]. Here we integrate this regulatory network with another that directly mediates protein stability by showing that the activities of DA1 peptidase are inhibited by BRI1- and BAK1-mediated phosphorylation. Through this regulation, BR-mediated signalling modulates DA1 substrate abundance, the extent and duration of cell proliferation during organ growth, and final organ size.

### Receptor kinases phosphorylate DA1 and modulate its peptidase activity

Receptor-like kinases (RLKs) can function as surface receptors that initiate signal transduction cascades through their extracellular domains by binding ligands, leading to the activation of their cytoplasmic kinase domains and subsequent phosphorylation of substrates. RLKs integrate diverse external signals; for example BAK1, a member of the SERK family, functions in concert with BRI1 [35,36] and FLS2 [37–39] to integrate BR- and pathogen-mediated signals [40]. We show that DA1 is another substrate of BRI1 and of BAK1 (Figure 4), extending the range of phosphorylation activities of both kinases and suggesting that DA1 family members may transduce signals from a wider range of RLKs through the association of BAK1 with these diverse RLKs. This conjecture is supported by the interaction of DA1 with the cytoplasmic kinase domain of the RLK TMK4 (which also interacts with BAK1 [25]), recently identified as a negative regulator of auxin biosynthesis [26].

Phosphorylation of DA1 at T367, S369 and T370 by BRI1 and BAK1 strongly reduced its peptidase activities and its repression of organ growth without influencing mono-ubiquitylation (Figure 5), which activates and inactivates DA1 peptidase [14,41]. BRI1 and BAK1 phosphorylate DA1 in a highly conserved domain near to the active site (Figure 4A). The *da1-1* negative interfering allele R358K and the *sod1-2* allele L362F that suppresses *da1-1* are adjacent to these phospho-sites [13,14], supporting the functional significance of this phospho-domain in modulating peptidase activity. As monoubiquitylation still occurred in the DA1-3D phospho-mimic protein, phosphorylation could modulate the peptidase activity of ubiquitylated forms of DA1. This is supported by the increased growth-repressive activity of non-phosphorylatable DA1-3A mutants (Figure 5C), suggesting a dynamic balance between phosphorylation and monubiquitylation may regulate peptidase activity. Mono-ubiquitylation of the DA1-3D phospho-mimic protein may contribute to a residual peptidase activity that was not detectable in enzymatic assays (Figure 5A) but was apparent by partial complementation of the *da1dar1* double mutant (Figure 5C). Currently it is not clear how this dual regulation interacts to control DA1 activity during growth. Figure 1 shows the proliferative zone occupies a larger proportion of the leaf at early stages of growth in the *da1-1* mutant. Although distributions of BR in developing leaves are not well characterised, the expression of BR biosynthetic enzymes is high in proliferating cells and reduced in leaf margins where growth is low [22] and BR promotes cell proliferation during leaf growth [18]. Therefore it is plausible that higher levels of BR promote cell proliferation during early stages of leaf growth in part by reducing the growth-repressive activities of DA1 by BRI1- and BAK1-mediated phosphorylation. Lower BR levels, for example at leaf margins, could increase DA1 activity, leading to reduced cell proliferation in these zones.

Regulatory interactions between ubiquitylation and phosphorylation is common and has previously been observed in the BR signalling pathway. Polyubiquitylation of BRI1 by the U-box E3 ligases PUB12 and PUB13 depends on its kinase activity and is required for endocytosis and control of protein levels [42,43]. BZR1 protein levels are modulated by the U-Box40 E3 ligase [44] and by sumoylation [45]. Cross-talk influencing DA1 activity is fundamentally different as it does not involve ubiquitin-mediated targeting or proteasomal degradation [14,44]. Instead, BL treatment and phosphorylation increased higher-MW forms of DA1 (Figure 5F) that contain DA1 homomeric forms. Reduced levels of high MW forms of DA1-3A and increased levels of high MW forms of DA1-3D indicate that these high MW complexes are less active forms of DA1. LIM and LIM-like domains comprising pairs of Zn finger motifs that characterise the DA1 family [13] confer specificity to protein-protein interactions, including dimerization [46]. DA1 family LIM domains may promote interactions in response to phosphorylation, and heterodimerization of functional redundant DA1, DAR1 and DAR2 proteins [47] may be a general mechanism controlling their activities.

### BR-mediated DA1 peptidase activity influences the levels of proteins involved in growth responses

Previous studies of BR-mediated responses have mainly focussed on transcriptional control mediated by a phospho-cascade terminating at members of BZR1 and BES1 families of transcription factors [48]. Our description of a direct role for BR signalling in the rapid modulation of levels of the DA1 substrates BB-Venus and UBP15-GFP (Figure 6) extends the scope of BR regulation of post-translational control of protein levels *via* the N-end rule pathway [49]. One mechanisml arising from these observations involves high BR levels maintaining the proliferative potential of cells by inhibiting DA1-mediated protein cleavage, while reduced BR levels re-programme the proteome by releasing peptidase inhibition, facilitating the transition to cell growth, endoreduplication and differentiation.

## STAR Methods

### Key Resources Table

| REAGENT or RESOURCE | SOURCE | IDENTIFIER |
| --- | --- | --- |
| **Antibodies** |  |  |
| HRP anti-FLAG | Sigma | Cat#A8592 |
| HRP anti-HA 3F10 | Sigma | Cat#12013819001 |
| HRP anti-GFP | Miltenyi | Cat#130-091-833 |
| mouse anti-tubulin | Sigma | Cat#T5168 |
| HRP rabbit anti-mouse | ThermoFisher | Cat#61-6520 |
| GFP-Trap-A beads | Chromotek | Cat#GTMA-20 |
| HA-magnetic beads | Pierce | Cat#88836 |
| FLAG-magnetic beads | Sigma | Cat#M8823-1ML |
| Rabbit anti Arabidopsis H+ATPase | Agrisera | Cat#AS07260 |
| HRP-mouse anti-rabbit | Sigma | Cat#MAB210P |
| Glutathione Sepharose 4B | Sigma | Cat#GE17-0756-01 |
| **Bacterial and Virus Strains** |  |  |
| E. coli DH5-alpha | NEB | Cat C287K |
| E. coli BL21 DE3 pLys/S | NEB | Cat C253OH |
| Agrobacterium tumefaciens GV3101 | [50] | N/A |
| **Biological Samples** |  |  |
| **Chemicals, Peptides, and Recombinant Proteins** |  |  |
| Kanamycin | Sigma | Cat#K1377 |
| Ampicillin sodium salt | Sigma | Cat#A0166 |
| Hygromycin | Sigma | Cat#H9773 |
| Spectinomycin | Sigma | Cat#S4014 |
| BASTA (glufosinate-ammonium) | Sigma | Cat#49677 |
| Murashige and Skoog plant growth medium | Duchefa | Cat#M0223 |
| epi-Brassinolide (BL) | Sigma | Cat#E1641 |
| Brassinazole (BRZ) | Sigma | Cat#SML1406 |
| Skim Milk Powder | Tesco | N/A |
| Benzonase | Sigma | Cat#E1014 |
| Yakult Macerozyme | Duchefa Biochemie | Cat#M8002 |
| Cellulase Onozuka R10 | Duchefa Biochemie | Cat#C8001 |
| PEG4000 | Sigma | Cat#P5413 |
| Tween-20 molecular biology grade | Sigma | Cat#P9416 |
| DDM N-dodecyl-b-D-maltoside | Sigma | Cat#850520P |
| EDTA-free Protease Inhibitor Cocktail tablet | Sigma | Cat#11873580001 |
| PhosSTOP phosphatase inhibitor tablet | Sigma | Cat#4906837001 |
| AESBF serine protease inhibitor | Sigma | Cat#A8456 |
| PVDF membrane 0.45um | ThermoFisher | Cat#88518 |
| IEF protein markers 3-10 | Serva | Cat#39212 |
| NativeMark gel protein markers | Life Technologies | Cat#LC0725 |
| SDS PAGE pre-stained protein markers | NEB | Cat#P7719S |
| Insta-blue protein stain | Expedeon | Cat#SKU:ISB1L |
| Novex Iso-electric focusing (IEF) pH 3-10 gel | Invitrogen | Cat#EC6655 |
| Novex Zoom tris-glycine 4-20% polyacrylamide gel | Invitrogen | Cat#EC6026 |
| NuPAGE 4-12% Bis-Tris SDS polyacrylamide gel | Invitrogen | Cat#NP0322 |
| Phos-TAG gel 6% | Fuji | Cat#192-17401 |
| SuperSignal West Femto HRP Substrate | ThermoFisher | Cat#34096 |
| SYBR green PCR master mix | ThermoFisher | Cat#4309155 |
| Qiagen RNEasy Kit | Qiagen | Cat#74904 |
| Qiagen maxiprep plasmid kits | Qiagen | Cat#12162 |
| GeneArt site-directed mutagenesis PLUS | Invitrogen | Cat#A14604 |
| Quant-iT protein assay kit | ThermoFisher | Cat#Q33210 |
| NuPAGE 4x LDS sample buffer | ThermoFisher | Cat#NP0007 |
| Novex Tris-Glycine 2x SDS sample buffer | ThermoFisher | Cat#LC2676 |
| Ubiquitin Human recombinant | Boston Biochem | Cat#U-100H |
| TEAB buffer 1M | Sigma | Cat#T7408 |
| Iodoacetamide | Sigma | Cat#I6125 |
| Acetonitrile | Sigma | Cat#27004 |
| Imidazole | Sigma | Cat#I5513 |
| Reduced glutathione | Sigma | Cat#G4251 |
| Acclaim PepMap C18 HPLC column | ThermoFisher | Cat#164567 |
| gamma <sup>32</sup>P ATP 6000Ci/mmol | Perkin Elmer | Cat#BLU002Z250 UC |
| **Deposited Data** |  |  |
| **Experimental Models: Cell Lines** |  |  |
| **Experimental Models: Organisms/Strains** |  |  |
| Arabidopsis thaliana Col-0 | Georg Redei, Univ. Missouri | N/A |
| Arabidopsis thaliana Col-0 bri1-301 | [28] | N/A |
| Arabidopsis thaliana Col-0 da1dar1bri1-301 | This Study |  |
| Arabidopsis thaliana Col-0 da1dar1 | [14] |  |
| Arabidopsis transgenics |  |  |
| pDA1::gDA1-mGFP in da1dar1 | This Study |  |
| pBRI1::BRI1-mGFP in Col-0 | Hannes Vanhaeren, VIB Gent | N/A |
| pDA1::gDA1-mGFP pBRI1::BRI1-mGFP | This Study |  |
| pDA1::gDA1-3D-mGFP in da1dar1 | This Study |  |
| pDA1::gDA1-3A-mGFP in da1dar1 | This Study |  |
| 35S::HA-DA1 in da1dar1 | This Study |  |
| 35S::HA-DA1-3D in da1dar1 | This Study |  |
| 35S::HA-DA1-3A in da1dar1 | This Study |  |
| pBB::BB-Venus in eod1-2 | This Study |  |
| pUBP15::UBP15-mGFP | [15] |  |
| **Oligonucleotides** |  |  |
| Primers are listed in Table S1 |  |  |
| **Recombinant DNA** |  |  |
| Bacterial Protein Expression |  |  |
| pETnT-FLAG-DA1-HIS | [14] |  |
| pETnT-FLAG-DA1-3D-HIS | This Study |  |
| pGEX4T2-GST-DA1 | [14] |  |
| pGEX4T2-GST-DA1-3A | This Study |  |
| pGEX4T2-GST-DAR1 | [14] |  |
| pGEX4T2-GST-DAR1-3A | This Study |  |
| pET24A-BAK1cd-HIS | This Study |  |
| pET24A-BAK1cd-HIS K317E kinase dead | This Study |  |
| pET24A-BRI1cd-HIS | This Study |  |
| pET24A-BRI1cd-HIS D1009N kinase dead | This Study |  |
| Yeast-2-Hybrid Interaction |  |  |
| pDEST GAD T7 DA1, DA1-3D, DA1-3A | This Study |  |
| pDEST GBK T7 DA1, DA1-3D, DA1-3A | This Study |  |
| Plant Protein Expression |  |  |
| pEARley 35S::3FLAG-DA1 | [14] |  |
| pEARley 35S::3FLAG-DA1-3A | This Study |  |
| pEARley 35S::3FLAG-DA1-3D | This Study |  |
| pB7 35S::3HA-DA1 | [14] |  |
| pB7 35S::3HA-DA1-3A | This Study |  |
| pB7 35S::3HA-DA1-3D | This Study |  |
| pEARley103 35S::DA1-GFP | This Study |  |
| pEARley 35S:3FLAG-UBP15 | [14] |  |
| pw1211 35S::BB-3FLAG | [14,51] |  |
| pw1211 35S::BRI1-3FLAG | This Study |  |
| pw1211 35S::BAK1-3FLAG | This Study |  |
| **Software and Algorithms** |  |  |
| Image J | NIH, USA | http://rsb.info.nih.gov/ij |
| Excel | Microsoft |  |
| Leaf Analysis Toolkit | [51] | (https://github.com/rosscarter3/leaf_analysis_toolkit) |
| Gaussian Projection | [51] | https://github.com/JIC-CSB/gauss_proj |
| R Statistical Analysis |  | https://www.r-project.org/ |
| Scaffold 4 |  | www.proteomsoftware.com |
| MaxQuant 1.6.0.16 |  | https://maxquant.org/ |
| Other |  |  |

### Resource Availability

Further information and requests for resources and reagents should be directed to and will be fulfilled by the Lead Contact Michael Bevan

#### Materials Availability

All plasmids generated in this study are available from the lead Contact without restriction.

#### Data and Code Availability

Code is available from Github as described in the Key Resources Table.

Phenotypic data tables are available from the Lead Contact.

### Experimental Model and Subject details

*Arabidopsis thaliana* accession Col-0 was used for transformation and as the source of T-DNA mutants. The *bri1-301* allele originated from an EMS-mutagenised population of Col-0 [28]. A transgenic Col-0 line (pAR169) expressing *pAtML1::mCitrine-RCI2A* was kindly provided by Dr Samantha Fox with the permission of the originator Dr Adrienne Roeder (Cornell University). This was crossed with the *da1-1* mutant [13] and plants with homozygous loci were identified. A pBRI1::BRI1-GFP line, kindly provided by Dr Hannes Vanhaeren (VIB Gent) was crossed to the *da1dar1* double mutant.

## Method Details

### Plant Materials and Growth Conditions

Arabidopsis plants were grown in Arabidopsis mix or half-strength Murashige and Skoog (1/2MS) medium plus 1% Bacto-agar at 22°C 16hr light / 8 hr dark cycles. For protoplast preparation 4-5 week old seedlings were grown in Arabidopsis mix at 22°C in 8 hr light / 16 hr dark cycles in an incubator. *bri1-301* lines were grown at either 22°C or the non-permissive temperature of 29°C.

### Protoplast Isolation and transfection

Protoplasts were isolated from *da1dar1* seedlings to reduce endogenous DA1 and DAR1 peptidase activities. The lower epidermis of leaves of 3-4 week old plants grown on soil was stripped using Scotch magic tape [52] and the remaining leaf tissue incubated with 1.5% cellulase Onozuka R10 and 0.4% macerozyme in W5 medium (20 mM KCl, 0.4M mannitol, 20 mM MES pH5.7) and incubated at 20°C with gentle shaking for 3 hrs. The digestate was filtered through 100 μm nylon mesh and centrifuged at 100 x g for 3 min. Protoplasts were gently re-suspended in cold W5 medium, incubated on ice for 30 min, centrifuged and resuspended in MMg solution (0.4M mannitol, 15mM MgCl_2_, 4 mM MES pH5.7) to 2 x 10^5^ protoplasts/ml. DNA (up to 20 μg in 20 μl) was added to 200 μl protoplasts, and 220 μl PEG/Ca solution was added (40% PEG 4000 in 0.2M mannitol, 100 mM Ca(NO_3_)_2_, pH 5.7). The protoplasts were incubated at room temperature for 30 min, diluted with 400 μl W5 solution, centrifuged, the supernatant completely removed, and the transfected protoplasts resuspended in 200 μl W5. This was dispensed into a 24 well microtitre plate and incubated at 20°C overnight in constant low light. Protoplasts were collected by centrifugation at 100x g for 2.5 minutes, and protein extracted as described below.

### Plasmid Construction and Plant Transformation

Plasmid constructs were made as described previously [14] using Gateway, Infusion or classical ligation methods. Arabidopsis transformation was carried out by floral dip of *Agrobacterium tumefaciens* GV3101 containing binary vector constructs. Stable transformants were selected on BASTA, hygromycin or kanamycin, and homozygous single copy insertion lines identified. The BRI1 cytoplasmic region used was amino acids 814-1196, and the BAK1 cytoplasmic region was amino acids 297-662.

### Mass spectrometry of phosphorylated DA1

GST-DA1, expressed from pGEX4T2, and BRI1-cd-HIS, expressed from pET28a, were co-transfected into *E. coli* BL21 pLys/S and selected on ampicillin and kanamycin. IPTG induction of protein synthesis was carried out at 28°C for 3 hrs, after which GST-DA1 was purified by Glutathione Sepharose 4B and eluted with 2x SDS sample buffer. The eluate was electrophoresed on 50 mM Phostag gels (6%) and bands with low mobility were excised. Gel slices were prepared according to standard procedures adapted from [53]. Gel slices were washed with 50 mM TEAB buffer pH8 (Sigma), incubated with 10 mM DTT for 30 min at 65 °C followed by incubation with 30 mM iodoacetamide (IAA) at room temperature in 50 mM TEAB. After washing and dehydration with acetonitrile, the gels were soaked with 50 mM TEAB containing 10 ng/μl Sequencing Grade Trypsin and incubated at 37 °C for 16 h. Peptides were extracted, and aliquots were analysed by nanoLC-MS/MS on an Orbitrap Fusion™ Tribrid™ Mass Spectrometer coupled to an UltiMate^®^ 3000 RSLCnano LC system. Analysis was performed using parallel CID and HCD fragmentation with fixed modification. The Mascot search results were imported into Scaffold 4 using identification probabilities of 99% for proteins and 95% for peptides. Recalibrated peaklists were generated with MaxQuant 1.6.0.16 [54] using the TAIR10_pep_20101214 *Arabidopsis thaliana* protein sequence database (www.arabidopsis.org) with 35,386 entries plus the Maxquant contaminants database. The final search was performed with the merged HCD and CID peaklists from MaxQuant using an in-house Mascot Server 2.4.1 on the same databases. Phosphorylation (STY), oxidation (M), deamidation (N/Q) and acetylation (protein N-term) were set as variable modifications.

### Yeast-two-Hybrid analyses of DA1 interactions

DA1, DA1-3D and DA1-3A were cloned into pDEST GAD and pDEST GBK to generate GAL4-AD and GAL4-BD clones respectively. These were co-transformed into the Matchmaker strains AH109 and Y187, and screens for interaction carried out on drop-out medium.

### Site-directed mutagenesis

This was carried using GeneArt site-directed mutagenesis PLUS using TOPO-D entry clones as substrates and primers described in Table S1.

### Seedling treatments

Hypocotyl elongation experiments used 1/2MS medium supplemented with 2.5 μM brassinazole (BRZ) or the equivalent volume of DMSO carrier solution. Seeds were stratified on 1/2MS medium at 4°C for 4 days, transferred to 16hr light for 24 hrs, transferred to plates containing DMSO or BRZ and grown vertically in the dark at 22°C or 29°C for 5 days. For *in vivo* phosphorylation analyses, 10-12 day old seedlings grown on 1/2MS plates were harvested, rinsed in 1/2MS liquid medium and incubated in 1/2MS medium with 2.5 μM BRZ for approximately 16 hrs. Seedlings were then rinsed in 1/2MS and then transferred to either 1/2MS + 1 μM BL or 1/2MS+ 2.5 μM BRZ for 1-2 hrs before harvest.

### Protein expression, purification and analysis

Proteins were extracted from seedlings (2ml ice-cold extraction buffer per 1 gm seedlings frozen and ground in LN2) or protoplasts (200 μl per transfection) using 50mM Tris HCl pH7.5, 150 mM NaCl, 10% v/v glycerol, 0.5 mM EDTA, 0.5% NP40, 1 mM AESBF and Roche protease inhibitor cocktail. For phosphoprotein analysis proteins were extracted in a protein extraction buffer supplemented with 1 x PhosStop. Extracts were sonicated for 2 x 30 secs, centrifuged at 4°C at 15,000g and the supernatant used. Proteins immuno-purified on beads were washed with 50mM Tris-HCl pH 7.5, 150 mM NaCl, 1 mM AESBF, protease inhibitor cocktail and 0.5% NP40. Microsomes were extracted from seedlings according to [55] using 100 mM Tris-HCl pH 7.5, 25% w/v sucrose, 5% v/v glycerol, 10 mM EDTA pH 8.0, 10 mM EGTA pH 8.0, 5 mM KCl, 1 mM DTT, 1 mM PMSF, 1 mM AESBF, Roche protease inhibitor cocktail and PhosStop. Seedlings were extracted (1gm seedlings with 2 ml of buffer), sonicated and centrifuged at 600 g for 3 mins. The supernatant was re-centrifuged at 600 g and the supernatant centrifuged again in 300 μl aliquots for 2 hrs at 21,000 g. Membrane pellets were resuspended in 150 μl wash buffer (20 mM Tris HCl pH 7.5, 5 mM EDTA, 5 mM EGTA, 1 mM AESBF, 1 mM PMSF, protease inhibitor cocktail and PhosStop) and re-centrifuged at 21,000 g for 45 mins. Pellets were frozen in LN2 and stored at −80°C. Protein levels were quantified using a Quant-IT protein analysis kit and a Qubit fluorometer. Microsome preparations for localising DA1 were carried out on 10-12 day old seedlings according to [14,56].

For *in vitro* kinase reactions, HIS-tagged BRI1-cd and BAK1-cd fusion proteins, and GST-tagged DA1 and DAR1 proteins were expressed in *E. coli* BL21, purified on Ni-NTA magnetic beads or glutathione beads and eluted with 300 mM imidazole or glutathione respectively. Three μg of each protein were incubated in a total of 30 μl 50mM Tris-HCl pH7.5, 10 mM MgCl_2_, 10 mM MnCl_2_, 1 mM DTT, 1 μM ATP and 183 kBq [^32^P]-γ-ATP. Reactions were incubated at 30°C for 30min with mixing at 1000 rpm and stopped with SDS sample buffer. Phosphorylation was visualised using autoradiography of CBB-stained SDS-polyacrylamide gels.

Immunopurification used magnetic beads coupled to anti-HA, anti-FLAG or anti-GFP. Beads were incubated at 4°C for 2-4 hrs and washed with TBS containing 0.5% NP40, 1 mM AESBF, Roche protease inhibitor cocktail and PhosStop. Proteins were eluted with sample buffer at 96°C for 6 minutes.

### *In vitro* ubiquitylation reactions

These were carried out as described [14].

### Protein gel analyses

Protein extracts were denatured at 90°C for 5 mins with NuPAGE LDS sample buffer and electrophoresed on 4-20% NuPAGE Bis-Tris SDS PAGE using MOPS-SDS running buffer. Native proteins were extracted from protoplasts in protein extraction buffer containing 1% n-docecyl-b-D-maltoside (DDM) instead of NP40, native PAGE sample buffer was added to 1X and 1/20v/v of 5%w/v Coomassie G-250 was then added. Samples were maintained at 4°C before electrophoresis at 4°C on NativePAGE Bis-Tris Novex 4-16% gels using 150v for 60 mins and 250v for 60 mins. Before transfer to PVDF membranes, native gels were washed in 100 ml 0.1% SDS for 15 mins. Two-dimensional gel analysis of purified microsomal proteins was carried out by extracting microsomal pellets in protein extraction buffer containing 1% Triton X-100 instead of NP40, sonication and centrifugation at 15,000 g for 10 mins. 20 μl of supernatant was added to 20μl 2 x Novex IEF sample buffer pH 3-10 and loaded onto Novex pH 3-10 IEF gels. These were electrophoresed at 4°C 100v for 1 hr, 200 v for 1 hr and 500v for 30 mins. IEF gels were soaked in 20% ethanol for 10 mins, each lane was carefully excised from the gel and incubated in 2 ml 2X Novex SDS Tris-glycine sample buffer + 0.5ml ethanol for 5 mins before transfer to 1X Novex SDS Tris-glycine sample buffer. The stained IEF strip was laid into a single well of a 4-20% Novex ZOOM Tris-glycine gel. Proteins on gels were transferred to PVDF membrane at 100v for 1 hr using Tris-glycine transfer buffer containing 10% methanol.

### Immunoblotting

After transfer PVDF membranes were rinsed in PBS and treated with 5% skim milk powder in PBS for 1hr with gentle shaking at room temperature. Membranes were then treated with antibodies as indicated for 2hrs at room temperature, washed in PBS at least 5 times, and then treated with either secondary antibody or Supersignal HRP reagent. Gels were then exposed to X-Ray film (Fuji R100).

### RNA analysis

RNA was isolated using Qiagen RNeasy plant kits and QuantiTECT reverse transcription kits were used. RT-qPCR was performed using SYBR Green Real-Time PCR mastermix on a Roche Lightcycler 480. Primer sequences used for RT-qPCR are in Table S1. Primer efficiencies and relative expression calculations were performed according to methods described by [57].

### Image analysis

The abaxial face of three representative first leaves of Col-0 or *da1-1* seedlings expressing the membrane-localised epidermal-specific mCitrine-RCI2A protein fusion was imaged from 5 das (days after stratification) with 20X objective using a Leica SP5 (II) or Leica TCS SP8X confocal scan head on a Leica DM6000 microscope. Argon laser excitation wavelength was 514nm and fluorescence emission between 520-600nm was collected. Data was analysed using the Leaf Analysis Toolkit software [51]. Gaussian projection was used on Z-stacks to identify fluorescence from the epidermal layer, to accommodate leaf curvature and to reduce bleed-through from underlying tissues. Segmentation used manual seeding by GIMP 2.8.16. Segmented image .csv files contained a unique identifier for each segmented cell, its location in the leaf, cell area, circularity cell perimeter and relative density.

To assess the locations of BRI1-eGFP and DA1-mCherry, *da1dar1* leaf protoplasts were transfected with 35S::DA1-mCherry and 35S::BRI1-GFP and imaged 16 hrs after transfection. A Leica SP5 was used for imaging transfected protoplasts. GFP was excited at 488nm and detected at 509nm, and mCherry was excited at 587nm and detected at 610nm.

Petals and cotyledons were imaged by high-resolution scanning (3600 dpi; Hewlett Packard Scanjet 4370) and areas calculated using ImageJ software.

### Quantification and Statistical Analysis

Data was collated and analysed using Excel or R. Heatmaps were created using the “heatmaps.py” script to generate .png files that were painted according to parameter values.

## Supplemental Information

Supplemental Information can be found online

## Acknowledgements

We thank Alex Jones for advice on phosphoprotein analysis, Samantha Fox for pAR169 and advice on imaging, Hannes Vanhaeren for the BRI1::BRI1-GFP transgenic line. Kim Findlay for scanning electron microscopy, and Matthew Hartley for image analysis software. RP and JD were supported by BBSRC-funded PhD studentships. MWB was supported by Biological and Biotechnological Sciences Research Council (BBSRC) Grant BB/K017225 and Institute Strategic Grants GRO (BB/J004588/1) and GEN (BB/P013511/1).

## Author Contributions

HD and MWB developed the research programme, HD, CS, JD, RP, RC, NMcK and MWB conducted experiments and analyses, and MWB wrote the paper with input from all authors.

