## Supplemental Information for "The Receptor Kinase BRI1 promotes cell proliferation in Arabidopsis by phosphorylation- mediated inhibition of the growth repressing peptidase DA1"

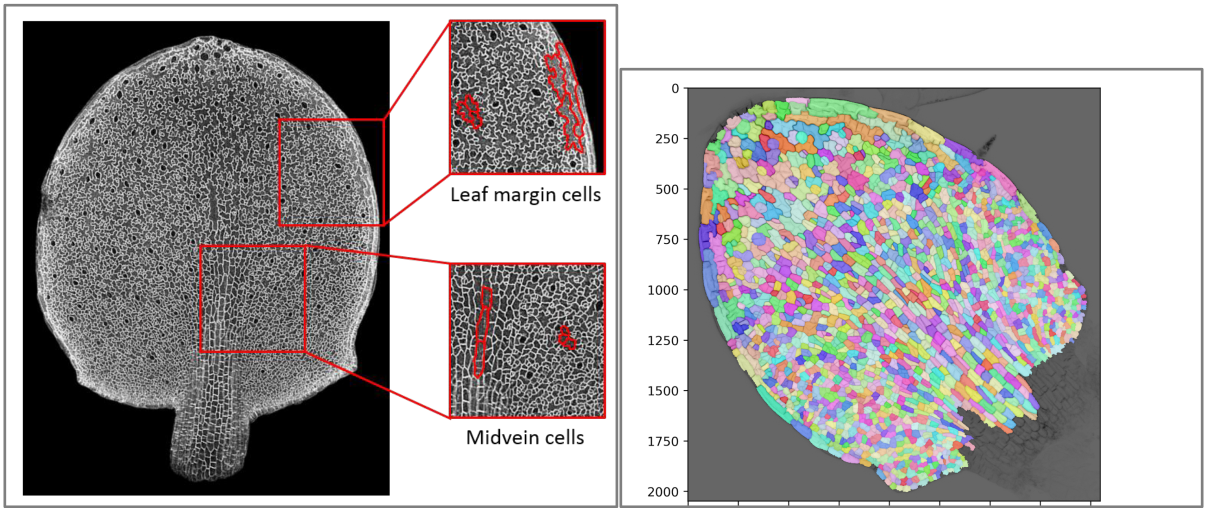


**Figure S1. An example of an imaged and segmented leaf. Related to Figure 1.**

The image on the left shows a typical confocal image of a whole pAR169 leaf that was used for segmentation (right image). At later stages of growth larger cells in the midvein and margins (compare shapes to adjacent pavement cells outlined in red) were excluded from segmentation as they confounded subsequent analyses. A typical segmented image of wt pAR169 at 7 das is shown on the right.

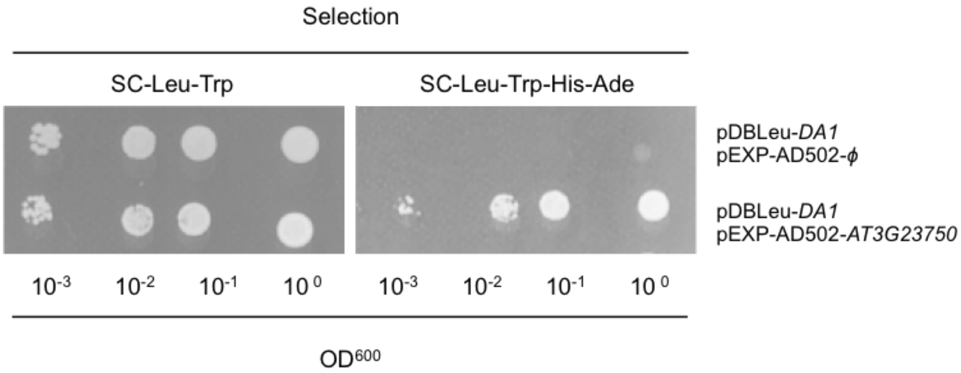


**Figure S2. Yeast- 2- hybrid screen identifies interactions between DA1 and the C-terminal region of the LRR TMK4. Related to Figure 3.**

The PJ69-4α yeast strain used in this screen is deficient for *LEU2* and *TRP1*, and had the *HIS3* and *ADE2* genes under the control of GAL4. The bait vector pDBleu contains the *LEU2* gene and expressed DA1. The prey vector pEXP-AD502 contained the *TRP1* gene and expressed an

Arabidopsis cDNA library (a gift from P. Wigge). Yeast co-expressing pDBLeu-DA1 and pEXP-AD-502-AT3G23750 (C-terminus of TMK4) were able to grow on SC$^{-Leu-Trp-His-Ade}$ medium, demonstrating a physical interaction. The negative control, DA1 with empty vector, was unable to grow on SC$^{-Leu-Trp-His-Ade}$ medium. Both bait and prey construct were able to grow on SC$^{-Leu-Trp}$ medium, demonstrating that they were being expressed.

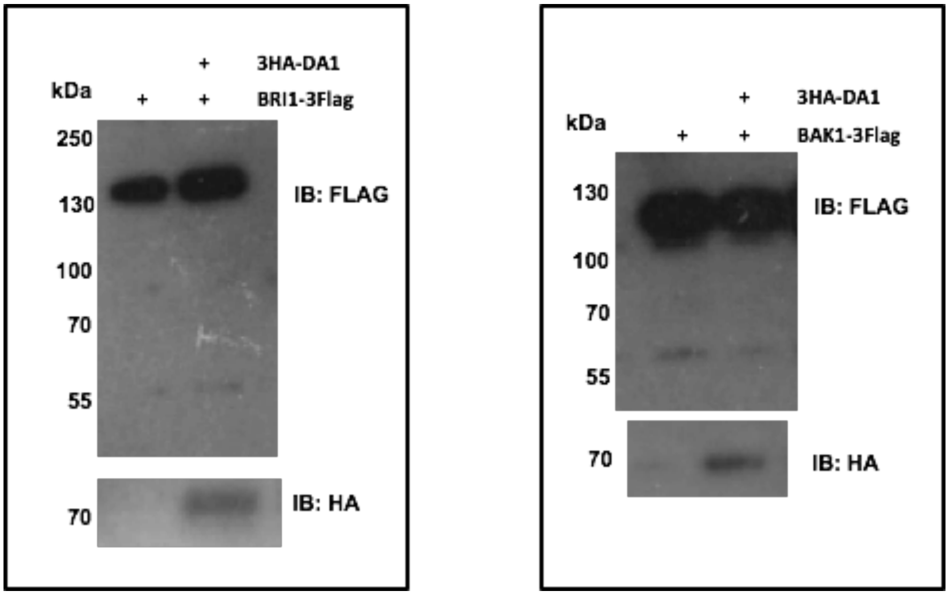


**Figure S3. DA1 does not cleave BAK1 or BRI1. Related to Figure 3.**

The immunoblots show BRI1-3FLAG or BAK1-3FLAG proteins co-expressed with HA-DA1 in *da1dar1* protoplasts. There was no evidence of HA-DA1 -mediated cleavage in these conditions.

A

B

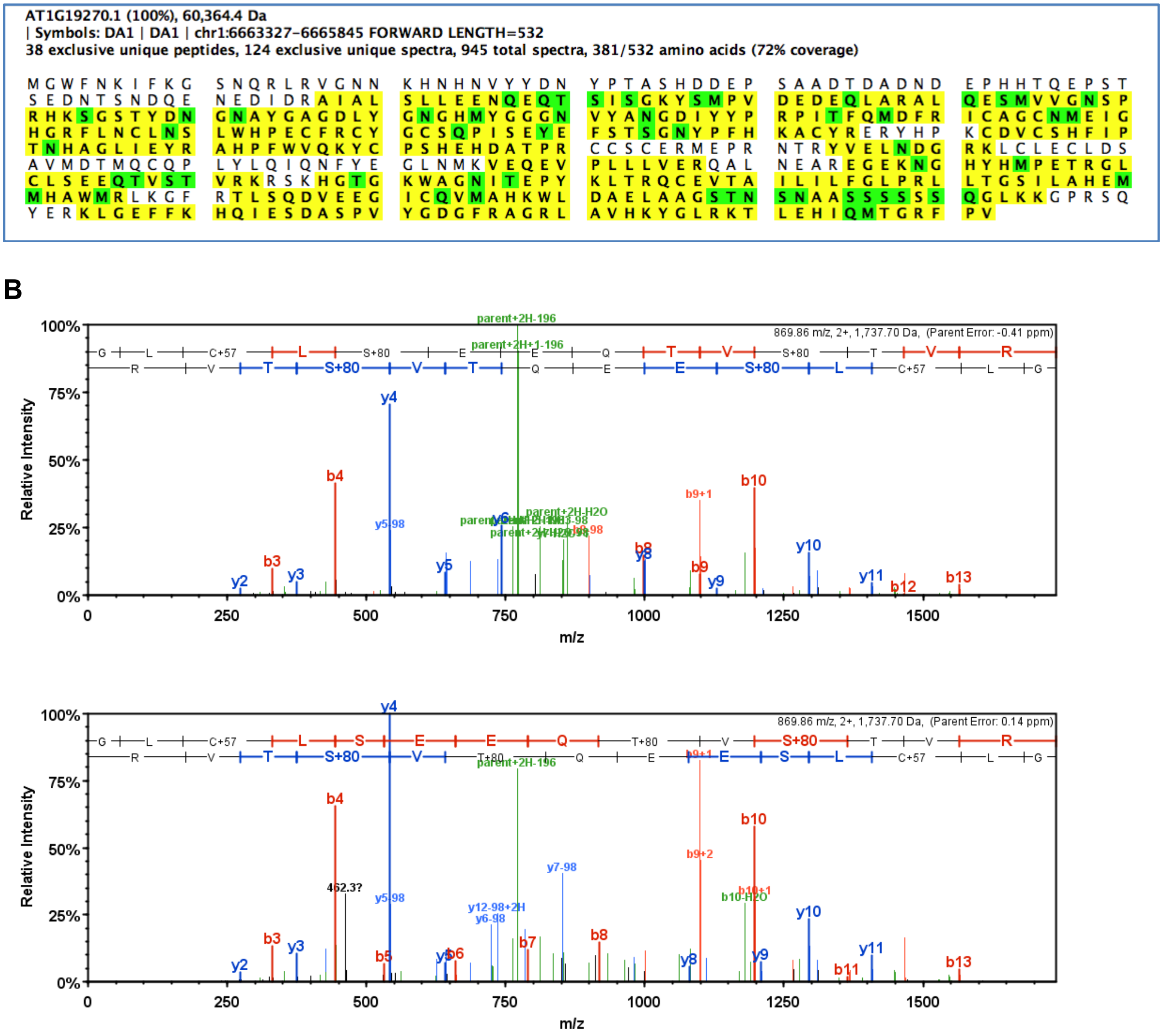


**Figure S4. Phospho-sites on DA1 from *E.coli* co-transfection with BRI1-HIS. Related to Figure 4.**

A. The diagram shows tryptic peptides (yellow highlights) identified by mass spectrometry mapped to the DA1 protein sequence. Modifications are highlighted in green. M oxidation; S,T phosphorylation; N,Q deamidation.

B. The upper panel is an example of mass spectra showing phosphorylation of serine 363 and 369, and the lower panel shows phosphorylation of threonine 367 and 370. Phosphorylation of serine 363 was less abundant than that measured for the other amino acids.

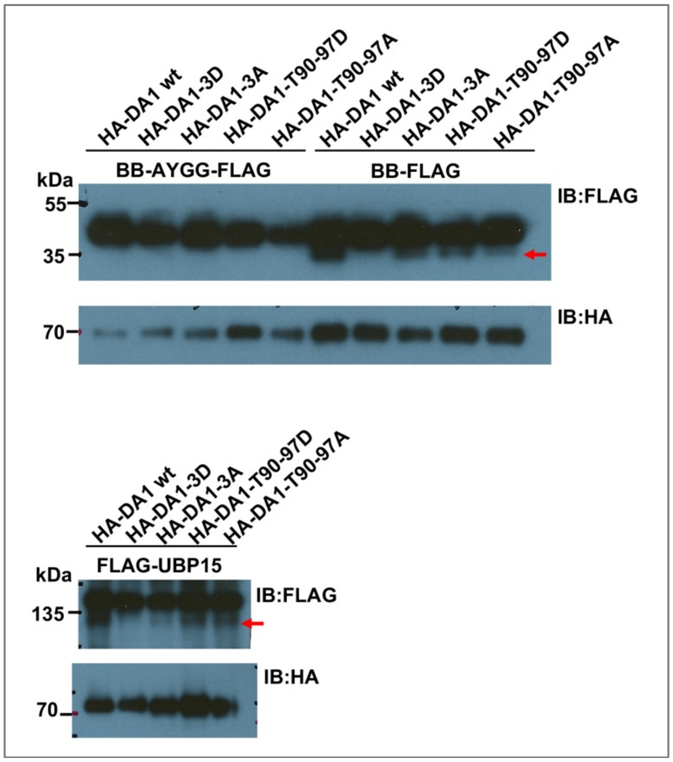


**Figure S5. Functional analysis of phospho-site mutations. Related to Figure 4.**

The panels are immune-blots of proteins expressed in *da1dar1* protoplasts. The top panels show co-expression of the non-cleaved BB-AYGG-FLAG protein (left) and BB-FLAG protein (right) together with wt HA-DA1 and different phospho-mutants. BB-AYGG-FLAG has the DA1 peptidase cleavage site NAYK mutated to NGGK. The lower panels are co-expression with the substrate FLAG-UBP15. HA-DA1-3D is T367D, S369D, T370D, HA-DA1-3A is T367A, S369A, T370A. HA-DA1-T90-S97D is T90D, S91D and S97D, and HA-DA1-T90-S97A is T90A, S91A and S97A. HA-DA1-3D was not able to cleave either BB-FLAG or FLAG-UBP15. All other phospho-site mutants were able to cleave both BB-FLAG and FLAG-UBP15.

A

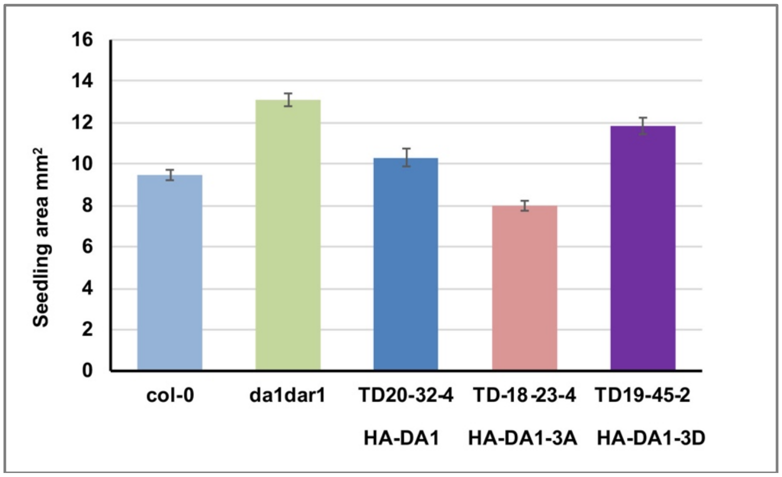


B

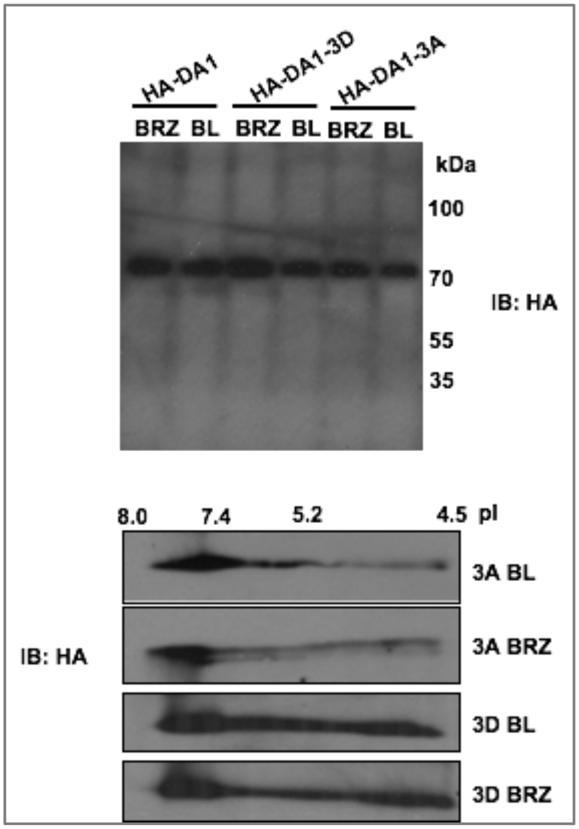


**Figure S6. Assessing microsome preparations from transgenic HA-DA1 wt, 3D and 3A seedlings. Related to Figure 4.**

Panel A. Complementation analysis of the transgenic lines HA-DA1, HA-DA1-3A, HA-DA1-3D used in microsome preparations.

Panel B. Immunoblot of microsome preparations. Each lane of the SDS-PAGE contains 20 μg of purified microsomes. A single band of HA-DA1 was seen.
The lower panels show immunoblots of 2D gels of microsome preparations from HA-DA1-3A and HA-DA1-3D seedlings treated with BRZ or BL. No alterations in pI in response to BL or BRZ were seen.

**Table S1 list of primers used, Related to STAR methods**

| Gene | Forward primer | Reverse primer | Function |
|---|---|---|---|
| **Cloning** | | | |
| DA1 infs Xba1 | AGATTACGCTTCTAGAATGGGTTG GTTTAACAAGATC | TAATTAACTCTCTAGATTAAACCG GGAATCTACCGG | Infusion N-HA |
| DA1-pro infs 3HA-DA1 | AGACCAAAGGGCTAT GTTAAAAATAAGCAGAAGAC | GTCCTCTCCAAATGA AATGCCTAAATAAAACTATC | Infusion N-HA |
| DA1 infs-cGFP Xho1 | GAGGACACGCTCGAG ATGGGTTGGTTTAACAAGATC | AGTCAGATCTACCAT AACCGGGAATCTACCGGTCATCTG | Infusion C-GFP |
| DA1 pro in-pEARley Xho1 | GAGGACACGCTCGAG AATTTTTGACAGAATAACAAC | GTTAAACCAACCCATAATGCCTAA ATAAAACTATCTG | Infusion |
| DA1 pro in-HADA1 no Xho1 R(×) | ATCGTAAGGGTACATAATGC CTAAATAAAA CTATCTG | TTGGAGAGGACACGCTCGAG AATTTTTGACAGAATAACAAC | Infusion |
| DA1 pro in DA1cGFP no Xho1 LR | ATCTTGTTAA ACCAACCCATAATGCCTAAATAAA ACTATC | GGAACATCGT AAGGGTACATAATGCCTAAATAAA ACTATC | Infusion |
| pDA1-Bpi1 F1, R1 | caGAAGACaaGGAGTCGTCGGATC CAATAGACGGCTAG | caGAAGACaaGTCaTCGTGT CCGAAGTTTT TTGG | Gateway |
| pDA1-Bpi1-F2,R2 | caGAAGACaatGACTTTTTAATTC ATTTCTCCTC | caGAAGACaaCATTATACGGCAAT GCAGCCTGCAAAATC | Gateway |
| DA1-CDS-Bpi1-F1 | caGAAGACaaAATGGGTTGGTTTA ACAAGATCTTTAAAG | caGAAGACaaGTCcTCATTT TCCTGGTCAT TCGATG | Gateway |
| DA1-CDS-Bpi1-R2 | caGAAGACaaCGAATTAACCGGGA ATCTACCGGTCATCTG | caGAAGACaaATGGATACGGCAAT GCAGCCTGCAAAATC | Gateway |

| | | | |
|---|---|---|---|
| gDA1-Bpi1-F3 | caGAAGACaaaACATGGTTTTGTTATCGGATTC | caGAAGACaaaACATGGTTTTGTTATCGGATTC | Gateway |
| DA1 in GST BamH1-Xho1 | GTTCCGCGTGGATCCATGGGTTGGTTTAACAAGATC | ATGCGGCCGCTCGAGTTAAACCGGGAATCTACCGG | Classical cloning |
| DA1 in pETnT BamH1-Xho1 | gattacgctggatccATGGGTTGGTTTAACAAGATC | gtggtggtgctcgagAACCGGGAATCTACCGGTCATC | Classical cloning |
| gDA1 pro-cds in ENTR- SmaI | GAATTCCTGCAGCCCGGGtattttgttgttgccttgcctacatac | AAGTTCTTCTCCTTTACTaaccgggaatctaccggtcatct | ENTR cloning |
| mGFP in ENTR- SmaI | GAATTCCTGCAGCCCGGGAGTAAAGGAGAAGAACTTTTCACTGGAGT | tttgggttTCACACGTGGTGGTGGTGGT | ENTR cloning |
| 3'DA1 in mGFP | CGTGTGAaacccaaatggacaaggtcttctac | TGGCGGCCGCTCTAGATTTCTTTCGCAAATACCATGGCTTTATCC | ENTR cloning |
| gDA1 pro-cds in ENTR- XbaI | TGGCGGCCGCTCTAGAaaccgggaatctaccggt | cccggttTTCgaacccaaatggacaaggtcttctact | ENTR cloning |
| BAK1 TOPO- | CACCATGGAACGAAGATTAATGATC | TCTTGGACCCGAGGGGTATTC | TOPO cloning |
| BAK1 infu-1 Xho1 | AGAGGACACGCTCGAG ATGGAACGAAGATTAATGATC | TATAATCCATCTCGAGTCTTGGACCCGAGGGGTATTC | Infusion |
| BAK1 in TOPO-EcoR1-Xho1 | GCCGCCCCCTTcaccATGGAACGAAGATTAATGATC | GGCGCGCCCACCCTTTCTTGGACCCGAGGGGTATTC | Classical cloning |
| BRI1-CD in-pET24a F EcoR1 - Xho1 | TCGCGGATCCGAATTCATGAGAGAGATGAGGAAGAG | GGTGGTGGTGCTCGAGTAATTTTCCTTCAGGAAC | Classical cloning |
| cBRI1 inTOPO F EcoR1 (lost) R Xho1 stop | CCCTTcaccGAATTCATGAAGACTTTTTCAAGCTTC | CCCACCCTTCTCGAGTCATAATTTTCCTTCAGGAAC | TOPO |
| gsGreen in pEARley103 Xho1 | TTTGGAGAGGACACGCTCGAGATGGTGAGCAAGGGCGAG | CTCCATTCATTGAACCGCCTCCACCCG | Classical cloning |

| | | | |
|---|---|---|---|
| EOD1 in gsGreen | AGGCGGTTCAATGAATGGAGATAATAGACCAGTGGAAGATG | TAATTAACTCTCTAGATGAACCGCCTCCACCCGGCTC | Classical cloning |
| **Site-directed Mutagenesis** | | | |
| DA1 T90-S97D | gacATAgacGGGAAAgACgacATGCCGGTGGATGAAGATG | gtcGTcTTTCCCgtcTATgtcgtcCTGTTCTTGATTCTC | P-site mutant |
| DA1 T90-S97A | gCAgcTATAgcCGGGAAAgcCgCGATGCCGGTGGATGAAG | cGgcTTTCCCGgcTATAgcTGcCTGTTCTTGATTCTCTTC | P-site mutant |
| DA1 T367A | CAAgCTGTTAGTACTGTAAGAAAG | ACTAACAGcTTGTTCTTCTGAAAG | P-site mutant |
| DA1 T367D | GAACAAgaTGTTAGTACTGTAAG | ACTAACAtcTTGTTCTTCTGAAAG | P-site mutant |
| DA1 T370A | GTTAGTgCTGTAAGAAAGCGATC | TCTTACAGcACTAACAGTTTG | P-site mutant |
| DA1 T370D | GTTAGTgaTGTAAGAAAGCGATC | TCTTACAtcACTAACAGTTTGTTC | P-site mutant |
| DA1 S369A | AAAGCATGGCgctGGAAAATGGG | TCCagcGCCATGCTTTGATCGC | P-site mutant |
| DA1 S369D | AAAGCATGGCgatGGAAAATGGG | TTTTCCatcGCCATGCTTTGATC | P-site mutant |
| DAR1 T385A T387A | CAAgcTGTCgCCACGGTGTTAAGG | CACCGTGGcGACAgcTTGTTCTTC | P-site mutant |
| DAR1 T388A | ATGATCgCAGAGCCTTGCAGGC | AGGCTCTGcGATCATGTCTATTAAC | P-site mutant |
| DAR1 T385D T387D | ATGATCgCAGAGCCTTGCAGGC | AGGCTCTGcGATCATGTCTATTAAC | P-site mutant |
| DAR1 T388D | CATGATCgacGAGCCTTGCAG | GGCTCgtcGATCATGTCTATTAAC | P-site mutant |
| BAK1 K317E | GGCCGTTgAAAGGCTAAAAGAGGAGC | AGCCTTTcAACGGCCACTAAAGTACC | Kinase-dead mutant |
| BRI1 D1009N | ACAGAaACATGAAATCCAGTAATGTG | ATTTCATGTtTCTGTGGATGATATGC | Kinase- dead mutant |
| **q-RT-PCR** | | | |
| *EXP8* | GATGGTCGCACACTCGTTAG | AGAGATGTGGATGATGGTTTGG | Gene expression |
| *SAUR-AC1* | ACTGTTGAGTAAATCTGAGGAAGAG | TGAGATGTGACTGTGAAGAACAA | Gene expression |
